# Cell-specific responses of *Anopheles gambiae* fat body to blood feeding and infection at a single nuclei resolution

**DOI:** 10.1101/2025.07.29.667437

**Authors:** Stephanie Serafim de Carvalho, Colton McNinch, Ana-Beatriz F. Barletta, Carolina Barillas-Mury

**Affiliations:** Laboratory of Malaria and Vector Research, National Institutes of Allergy and Infectious Diseases, National Institutes of Health; Rockville, Maryland, 20852, USA; Bioinformatics and Computational Biosciences Branch, Office of Cyber Infrastructure and Computational Biology, National Institute of Allergy and Infectious Diseases, National Institutes of Health; Bethesda, Maryland, 20892, USA

## Abstract

The mosquito fat body plays key roles in metabolism and immunity, yet its cellular diversity and functional specialization remain unclear. We characterized the *Anopheles gambiae* fat body and associated cells, examining their responses to blood feeding, bacterial infection, and immune priming following *Plasmodium berghei* infection, at single-cell resolution. We analyzed 97,650 nuclei from the female mosquito’s abdominal body wall and identified seven major cell types. Fat body trophocytes were the most abundant (∼85% of cells), while sessile hemocytes represented 7.4% of cells. Trophocytes consisted of five subpopulations, including basal (T1, T2), metabolic-enriched (T3), immune-responsive (T4), and a vitellogenic population (T5) exclusive to blood-fed females. T4 trophocytes exhibited constitutive expression of immune genes, while multiple cell types, including other trophocytes, hemocytes, and epidermal epithelial cells, responded to a systemic bacterial challenge. Oenocytes (1.1% of cells) induced the expression of enzymes involved in the biosynthesis of lipids in response to immune priming. Blood feeding triggered massive transcriptomic changes, with a strong induction of vitellogenin and multiple genes involved in DNA replication, consistent with trophocyte endoreplication and metabolic reprogramming. Interestingly, vitellogenin mRNA was expressed only in the first layer of trophocytes facing the hemolymph and had an apical subcellular localization. These findings provide a high-resolution atlas of fat body and associated cells, revealing specialized roles in immunity and reproduction and offering insights into how mosquitoes coordinate metabolic and immune functions at the cellular level.

## Introduction

Mosquito-borne diseases continue to impose a significant global health burden, with *Anopheles gambiae* as the primary African vector of *Plasmodium falciparum* malaria, the most severe form of the disease in humans. Female mosquitoes become infected when they ingest blood from an infected person containing gametocytes. The parasite undergoes a series of developmental stages in the vector, it multiplies, and is transmitted when the mosquito feeds on a new host. Blood feeding is an important physiological event in mosquitoes, essential for reproduction and egg development, and triggers activation of complex hormonal and nutrient-derived signaling pathways that regulate blood digestion in the midgut, yolk protein precursor (YPP) synthesis in the fat body during vitellogenesis, and nutrient uptake by developing oocytes in the ovaries. [1, 2].

The mosquito fat body is a multifunctional organ analogous to both the adipose tissue and liver in mammals. It is the primary site of nutrient storage and energy metabolism [3], contributes to the systemic immune response by producing antimicrobial peptides and participating in pathogen melanization [4], facilitates detoxification of metabolic waste and xenobiotics [5, 6], and is a major site of hemolymph protein synthesis [3]. It also plays a central role in vitellogenesis by synthesizing and secreting yolk proteins into the hemolymph that are transported to the ovaries to support egg maturation [1]. Structurally, the abdominal fat body consists of flattened lobes loosely attached to the abdominal integument that project towards the visceral organs and are in direct contact with the hemolymph [7, 8]. The integument, the outermost layer of the mosquito body, is formed by an outer layer of cuticle that is synthesized by a single layer of epidermal cells supported by a basal lamina [9]. The fat body lobes vary in size and are permeated by the respiratory system, which consists of a network of tracheoles [9]. Based on their anatomical position, the abdominal fat body can be described as ventral, lateral, or dorsal.

The abdominal body wall contains the fat body and other associated cells. Trophocytes are the predominant cell type within the fat body [3]. These large polyploid cells form the fat body lobes and accumulate nutrient reserves in the cytoplasm in the form of lipid droplets and glycogen granules [7, 8, 10–12]. Oenocytes are rounded cells derived from ectodermal tissue, that can be individual or organized in clusters [7, 13] and contribute to hydrocarbon synthesis for the cuticle [14], detoxification [15, 16], sex pheromone production [13, 17], and immune priming [18]. Immune priming in *An. gambiae* is mediated by the systemic release of a hemocyte differentiation factor (HDF) that consists of a complex formed by lipoxin A4 bound to Evokin, a lipid carrier [18–20]. This response is triggered when the microbiota comes in contact with the midgut epithelia during *Plasmodium* ookinete invasion of the midgut [20, 21]. HDF release increases the proportion of circulating granulocytes and enhances the immune response to subsequent infections [19–21]. Oenocytes express the double peroxidase enzyme (DBLOX), which mediates HDF synthesis, and their numbers increase in primed mosquitoes [18].

Other associated fat body cells include pericardial cells, hemocytes, and neurons. Pericardial cells (PCs) are located in the dorsal body wall along the mosquito heart [9]. They flank each ostium, a strategic site of high flow as hemolymph enters the heart to be pumped out and distributed throughout the mosquito hemocoel [9, 22, 23]. PCs are immunocompetent cells that sense pathogens in the hemolymph and are also involved in osmoregulation and detoxification [9, 22, 24, 25]. Hemocytes are immune cells that circulate in the hemolymph but are also often found associated with the surface of different tissues, including the fat body. For example, periosteal hemocytes aggregate on the heart surface and can phagocytize pathogens present in the hemolymph [22]. Mosquitoes have a ventral neural cord spanning the thorax and abdomen, as part of the nervous system that coordinates sensory and locomotor signals between the brain and the body [9].

Despite the central role of the fat body and its associated tissues in mosquito physiology and immunity, the cellular composition and cell-type-specific responses to blood feeding and infection remain poorly defined. In this study, we used bulk and single-nucleus RNA sequencing (snRNA-seq) to systematically characterize the abdominal fat body of *An. gambiae* adult females. By integrating molecular profiling with morphological analyses, we identified and classified the major cell types within the fat body and adjacent tissues. Furthermore, we investigated how these cell populations transcriptionally respond to physiological stimuli such as blood feeding, as well as bacterial and *Plasmodium* infection. Our findings provide a comprehensive cell atlas of the mosquito fat body and uncover dynamic transcriptional programs that underlie its diverse functional roles in metabolism, reproduction, and immunity.

## Results

### Single-nuclei sequencing reveals 12 cell clusters in the abdominal body wall of *An. Gambiae* female mosquitoes

A robust single-nucleus isolation protocol was established, yielding 700 to 1200 nuclei per microliter. The 10x Chromium technology was employed to perform 3’ snRNA-seq on pooled nuclei samples collected from the abdominal wall of healthy *An. gambiae* females fed on sugar or blood, as well as from females systemically challenged with bacteria or primed by infecting them with *Plasmodium berghei* (Figure 1A). Samples from females challenged systemically with a mixture of *Escherichia coli* and *Micrococcus luteus* were analyzed 4 hours post-injection, while the priming response was evaluated six days after females were fed on *P. berghei*-infected mice. The response of fat body cells to blood feeding was investigated 24 hours after ingestion of a blood meal.

**Figure 1:**
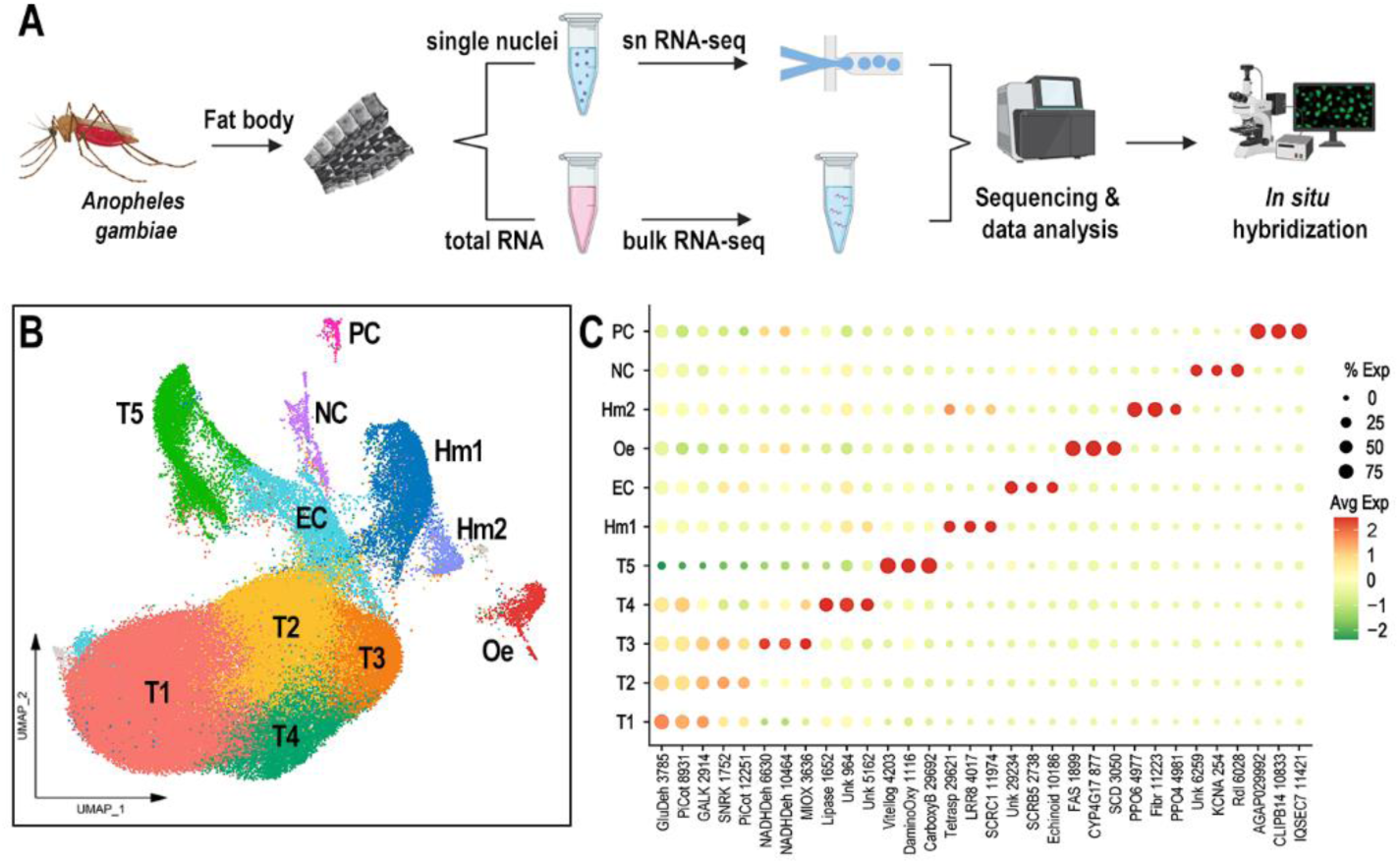
Molecular identification of *Anopheles gambiae* fat body cell types. (A) Schematic overview of the experimental workflow. (B) UMAP (Uniform Manifold Approximation and Projection) visualization of 97,650 nuclei isolated from *An. gambiae* body wall, colored by cluster identity. (C) Dot plot displaying three representative marker genes per cluster. Dot size indicates the percentage of cells within each cluster expressing the gene, while dot color represents the average expression level. Gene names were abbreviated, with numerical identifiers corresponding to VectorBase AGAP accession numbers. Cell type annotations: T1-T5 = Trophocytes, Oe = oenocytes, Hm1 and Hm2 = Hemocytes, EC = Epithelial cells, NC = Neural cord, and PC = Pericardial cells.

A total of 97,650 high-quality individual fat body nuclei were obtained from all samples combined. More than 2×10^9^ total sequence reads from the fat body nuclei were acquired, with a median of 949 transcripts and 560 genes per nucleus, and a transcriptome mapping ratio ranging from 57.6% to 76.6%. To ensure data quality, cutoffs for the number of counts and features, as well as the percentage of mitochondrial and ribosomal content, were set for each sample (Supplementary Figure 1). The datasets were integrated after filtering, resulting in 12 unsupervised clusters visualized using a Uniform Manifold Approximation and Projection (UMAP) (Figure 1B). Specific gene markers for each cluster were identified (Avg log2FC > 0.5, P val adj < 0.05), and candidate markers were selected for validation by *in situ* hybridization (ISH) (Supplementary Table 1, Figure 1C, Figure 2). One cluster had fewer than 300 cells after integrating all samples (colored in gray in the UMAP, Figure 1B) and could not be validated through ISH; therefore, it was not considered a relevant cell cluster and will not be further discussed.

**Figure 2:**
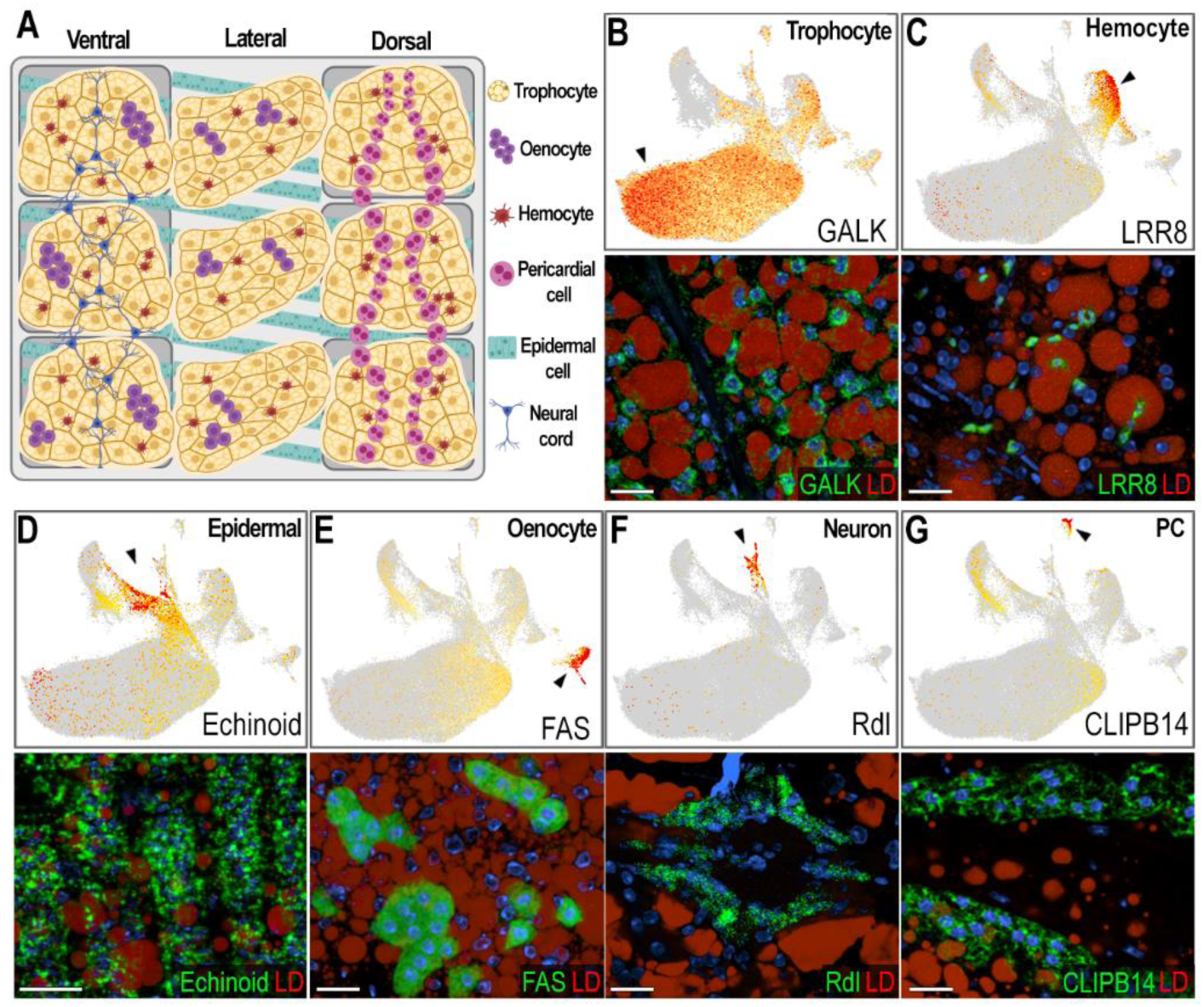
Validation and characterization of abdominal fat body cell types in female *Anopheles gambiae*. (A) Schematic illustrating the morphology and spatial distribution of abdominal fat body cell types. (B-G, top) Feature plots showing expression levels of marker genes for each cell type. Cells with log-normalized expression below 1 were shown in grey. Expression levels are shown in a yellow-to-orange gradient. (B-G, bottom) RNA *in situ* hybridization in sugar-fed females confirming marker gene expression in corresponding cell types. Marker transcripts are shown in green, lipid droplets (LD) in red, and nuclei in blue. Scale bars: 20µm. (B) Galactokinase (GALK; AGAP002914), a marker of sugar-fed trophocytes – clusters T1-T4. (C) LRR8 (AGAP004017), a marker of hemocytes of granulocyte lineage – cluster Hm1. (D) Echinoid (AGAP010186), a marker of epidermal cells – cluster EC. (E) Fatty acid synthase (FAS; AGAP001899), a marker of oenocytes – cluster Oe. (F) GABA-gated chloride channel (Rdl; AGAP006028), a marker of neurons that are part of the neural cord – cluster NC. (G) CLIPB14 (AGAP010833), marker of pericardial cells – cluster PC. Created in BioRender. Serafim de Carvalho, S. (2025) https://BioRender.com/ovwpy85.

### Identification and validation of the different cell types corresponding to the abdominal body wall clusters

Based on gene markers from each cluster and experimental validation through ISH, we identified eleven clusters corresponding to fat body trophocytes and other cells associated with the abdominal wall. We identified a total of seven different cell types associated with the abdominal body wall of female mosquitoes: fat body trophocytes with five subpopulations (T1, T2, T3, T4, T5), two types of hemocytes (Hm1, Hm2), epidermal cells (EC), oenocytes (Oe), neurons (NC), and pericardial cells (PC) (Figure 1B-C, Figure 2, Supplementary Table 1).

Trophocytes were the most abundant cell type, comprising 85.2% of the cells in the dataset (Supplementary Table 2). They are large polymorphic cells with cytoplasm filled with lipid droplets, organized in multiple layers that form the fat body lobes associated with the different segments of the abdominal body wall (Figure 2A-B). Five distinct trophocyte transcriptional profiles (T1, T2, T3, T4, T5) were observed, with Galactokinase (GALK - AGAP002914) serving as a general marker gene expressed in all trophocyte clusters (Figure 1C, Figure 2B). Surprisingly, hemocytes (Hm1, Hm2) are the second most abundant cell type, representing 7.4% of the cells (Supplementary Table 2). Some are associated with the hemolymph surface of the fat body, while others permeate the trophocyte layers in within the lobes (Figure 2A, Supplementary Figure 2). Hemocytes of the granulocyte (Hm1) lineage were the most prevalent (86.9% of hemocytes) (Figure 2C, Supplementary Figure 2), identified using Leucine-rich-repeat protein 8 (LRR8 - AGAP004017) as a marker, while Prophenoloxidase 4 (PPO4 - AGAP004981) expression identified oenocytoids (Hm2), representing 13.1% of hemocytes (Supplementary Figure 2). Epidermal cells (EC) are closely associated with the cuticle (Figure 2A, D), have small lipid droplets in their cytoplasm, and are organized in stripes (Figure 2D). They represent 3.26% of the isolated cells (Supplementary Table 2), and expression of Echinoid (AGAP010186) confirmed their identity in ISH.

The abdominal wall of adult females has three main regions: ventral, lateral, and dorsal, with significant differences in cell types associated with these cuticular regions (Figure 2A). Oenocytes (Oe) are present in the ventral and lateral regions, either as individual cells, or organized as cell clusters embedded in the abdominal fat body that lack lipid droplets in their cytoplasm (Figure 2A, E; Supplementary Figure 3A-B). They correspond to 1.12% of the cells (Supplementary Table 2), with fatty acid synthase (FAS - AGAP001899) as a specific gene marker (Figures 1C, 2E). Neurons from the neural cord (NC) were observed in the ventral region (Figure 2A, F), representing 0.84% of the cells, and expression of the resistance to Dieldrin gene (Rdl - AGAP006028), a GABA-gated chloride channel subunit, confirmed their identity by ISH (Figures 1C, 2F). Finally, pericardial cells (PC) (Figure 2G) are present in the dorsal region along the heart, representing 0.41% of cells, and express the Clip-domain Serine Protease B14 (CLIPB14-AGAP010833) as a specific marker (Figures 1C, 2A). The morphology and transcriptional characterization of the different cell types associated with the abdominal wall revealed a complex organization in the lateral, ventral, and dorsal regions.

### Oenocytes are the main cell type involved in immune priming

We performed bulk and snRNA-seq on mosquitoes challenged with *P. berghei* to assess the transcriptional and cell-specific responses to immune priming in cells associated with the mosquito body wall. Bulk RNA-seq revealed 45 upregulated genes and 1 downregulated gene (logFC > 0.5, FDR < 0.05) in the body wall of primed mosquitoes (Figure 3A, Supplementary Table 3) six days after *Plasmodium* infection. Notably, 42.2% of upregulated genes were markers of the oenocyte cluster (Figure 3A, genes with asterisk), with 37.7% previously reported as expressed or enriched in oenocytes [14, 15]. Most of these genes are related to fatty acid metabolism, including propionyl-CoA synthase (AGAP001473), fatty acid synthases (FAS - AGAP001899, AGAP008468, AGAP028049), fatty acid elongases (AGAP005512, AGAP007264, AGAP003195, AGAP003197), fatty acyl-CoA reductases (FAR2 - AGAP004784, AGAP005986, AGAP005984, AGAP005985), stearoyl-CoA desaturases (SCD - AGAP003050, AGAP003051), and cytochrome P450 enzyme (CYP4G17 - AGAP000877).

**Figure 3:**
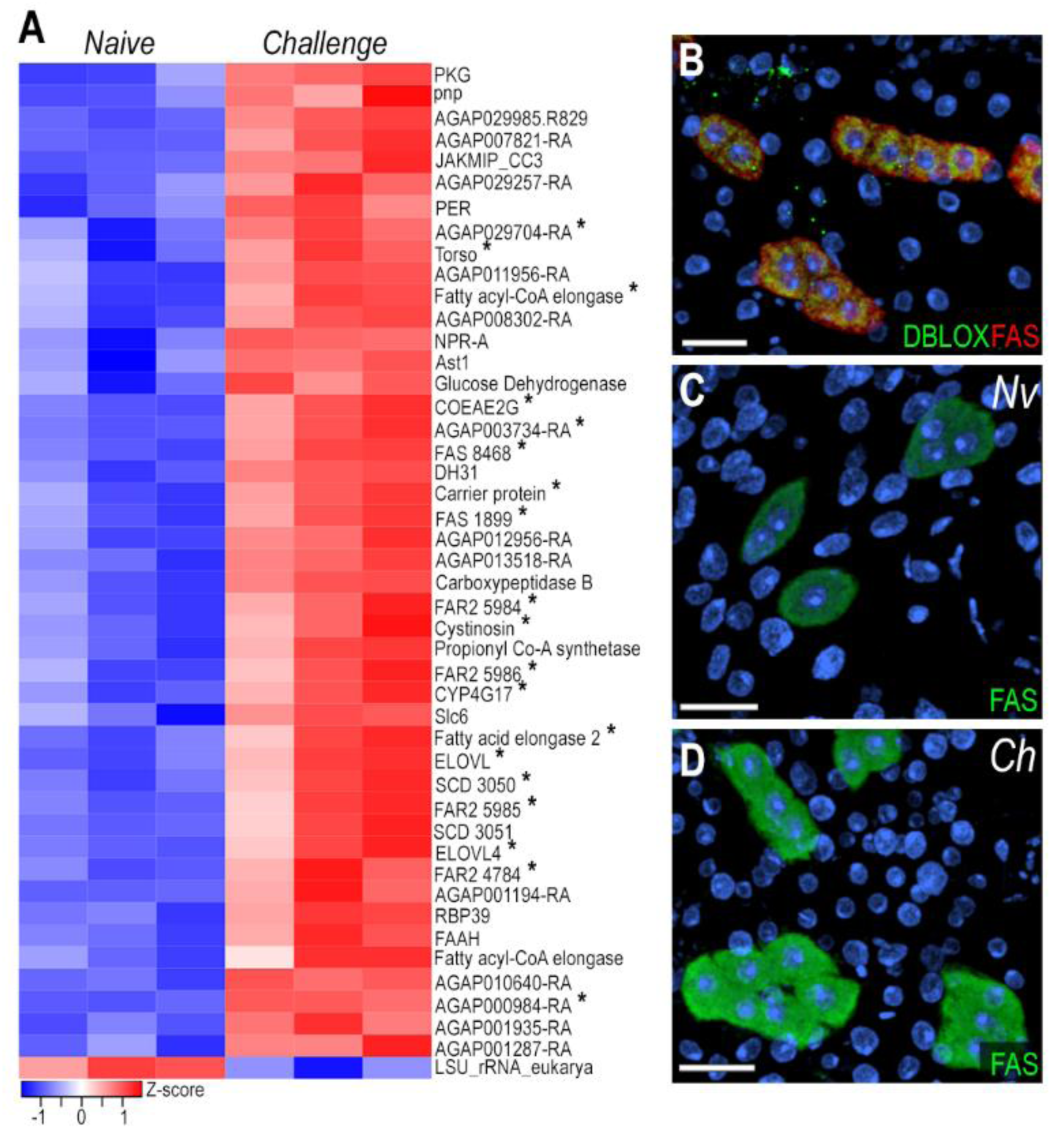
Oenocytes induce lipid synthesis genes following immune priming by *Plasmodium berghei* challenge. (A) Heatmap of differentially expressed genes from bulk RNA-seq analysis. Asterisk (*) indicate genes that are upregulated following *P. berghei* challenge and are also markers of oenocytes. (B) RNA *in situ* hybridization of DBLOX (AGAP008350) and FAS (AGAP001899) in oenocytes on the lateral abdominal body wall under naïve condition. DBLOX in shown in green, FAS in red, and nuclei in blue. (C,D) RNA *in situ* hybridization of FAS in oenocytes on the lateral abdominal body wall under naïve (C) and *P. berghei-* challenge (D) conditions. FAS is shown in green and nuclei in blue. Scale bars: 20µm.

Fatty acid synthase (FAS, AGAP001899) mRNA expression has strong and specific expression in the oenocyte clusters present in the ventral and lateral abdominal body wall (Figure 2E, Supplementary Figure 3A), but is absent in the dorsal region (Supplementary Figure 3B). We also stained the abdominal wall tissue with a specific probe to detect DBLOX (AGAP008350), an enzyme involved in lipoxin A4 synthesis, confirming the identity of oenocytes as FAS+ and DBLOX+ cells that lack lipid droplets and are individual or organized in clusters (Figure 3B, Supplementary Figure 3C-F). Furthermore, we confirmed by ISH that FAS expression is highly induced in response to priming in oenocytes present in the lateral and ventral regions of the abdominal wall (Figure. 3C-D, Supplementary Figure 3C-F). However, we could not detect a difference in DBLOX expression levels by ISH (Supplementary Figure 3C-D). Taken together, these observations indicate that oenocytes express many key enzymes involved in lipid biosynthesis and activate their transcription in response to immune priming.

Circulating granulocytes are the final effectors of the immune priming response. To investigate the response of granulocytes associated with fat body tissue, we performed differential gene expression analysis on the granulocyte cluster (Hm1) between naïve and challenged mosquitoes. This analysis revealed 75 upregulated genes and 135 downregulated genes (Avg log2FC > 0.5, P val adj < 0.05) (Supplementary Table 4). Some upregulated genes are related to extracellular matrix and adhesion, such as chondroitin sulfate synthase (AGAP001010) and matrix metalloprotease 2 (AGAP011870). Additionally, genes related to cholesterol synthesis were upregulated, including progestin and adipoQ receptor family member 3 (AGAP000144), Patatin-like phospholipase domain containing 5 (AGAP011925), and long-chain acyl-CoA synthetase (AGAP011603) (Supplementary Table 4). Among the downregulated genes, 18% are involved in the oxidative phosphorylation pathway, and 10% in carbohydrate metabolism (Supplementary Table 4). Although we observed this change in transcriptional response, we did not observe a significant difference in the number of body wall-associated granulocytes between naïve and primed mosquitoes.

### Functional characterization of fat body trophocyte subpopulations

Four trophocyte clusters (T1, T2, T3, and T4) were present in all sugar-fed samples analyzed, while an additional cluster (T5) appeared only in blood-fed females. In general, the transcriptional differences between trophocytes clusters are more subtle and unique markers cannot be used to identify specific subpopulations. T1 and T2 lack unique specific markers (Figure 1C), whereas clusters T3, T4, and T5 express some markers at higher levels and in a higher proportion of cells than other subpopulations (Figure 1C, Supplementary Table 1). Most markers in cluster T3 are related to protein translation and metabolic pathways such as glycolysis, the tricarboxylic acid cycle, and oxidative phosphorylation (Supplementary Table 1). In contrast, markers for cluster T4 are associated with immunity, melanization, or endoplasmic reticulum (ER)-related proteins (Supplementary Table 1). The expression of different functional classes of marker genes suggests that they represent distinct trophocyte subpopulations that perform specialized functions in the abdominal fat body.

To further characterize transcriptional differences between trophocyte clusters in sugar-fed mosquitoes, we performed ISH using galactokinase (GALK - AGAP002914) as a general trophocyte marker with higher expression in the T1 cluster; inositol oxygenase (MIOX - AGAP003636) as a marker enriched in T3, and lipase (AGAP001652) in the T4 trophocyte cluster (Figure 4A). Galactokinase was expressed in all trophocytes of sugar-fed females, with heterogeneity in the levels of expression (Figure 4B-C). Among the GALK+ trophocytes, some also expressed high levels of MIOX (GALK^+^, MIOX^h^) (Figure 4C-D), others expressed GALK and high levels of the lipase (AGAP001652) marker (GALK^+^, Lipase^h^) (Figure 4C, E); while a few trophocytes had strong expression of both markers (MIOX^h^, Lipase^h^) (Figure 4F, white arrow). These findings provide direct evidence that although trophocytes have similar morphology there is heterogeneity in the combination of markers they expressed, indicative of some level of specialization.

**Figure 4:**
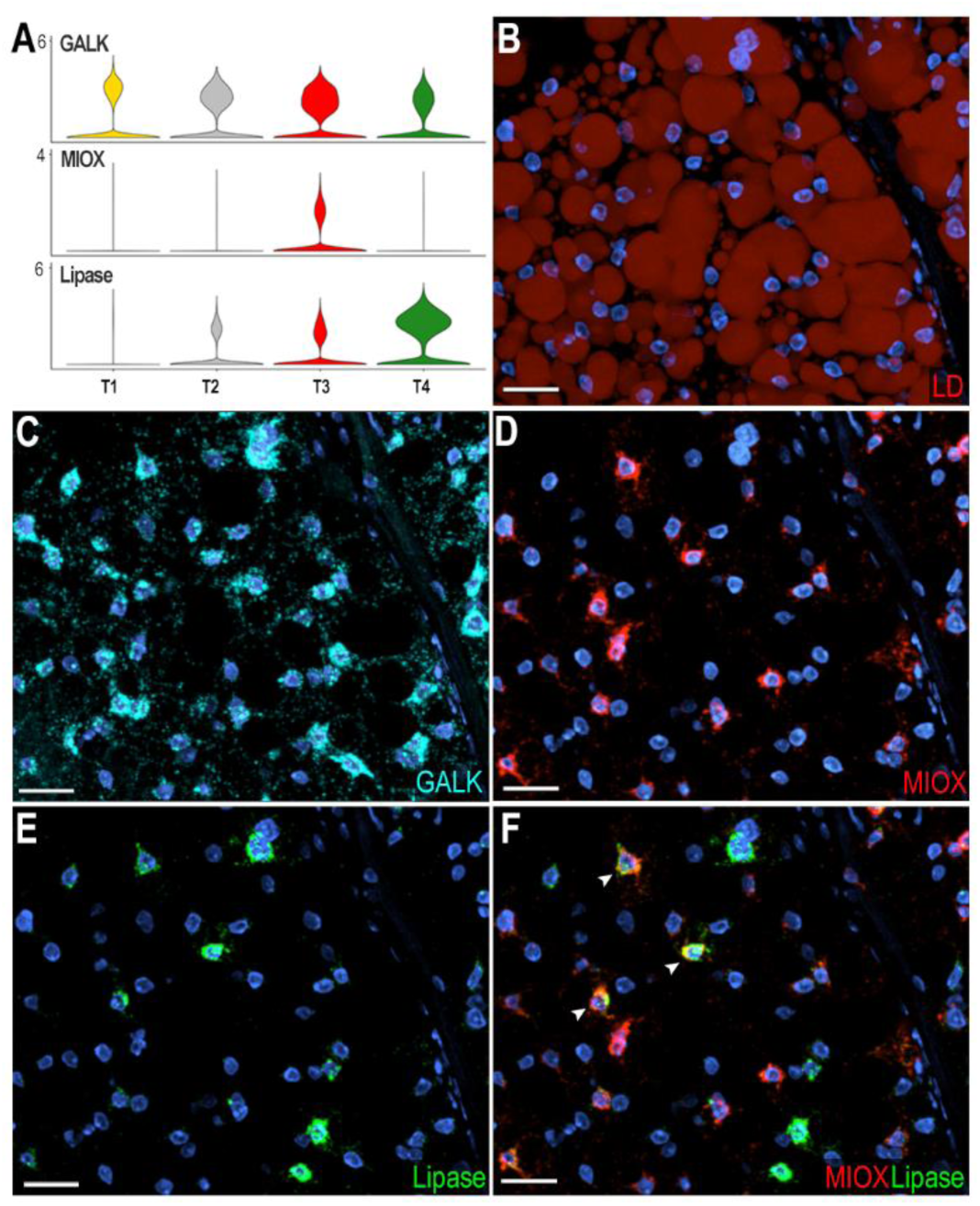
Characterization of trophocytes subpopulations in sugar-fed mosquitoes. (A) Violin plots showing expression of Galactokinase (GALK; AGAP002914), Inositol oxygenase (MIOX; AGAP003636), and Lipase (AGAP001652) in trophocytes. (B) Lipid staining of sugar-fed mosquito fat body showing lipid droplets (LD) distributed throughout the trophocyte cytoplasm. LD are stained in red. (D - G) RNA *in situ* hybridization validating trophocyte subpopulation markers: (D) GALK, marker of the T1 trophocyte subpopulation. (E) MIOX, marker of the T3 trophocyte subpopulation. (F) Lipase, marker of T4 trophocyte subpopulation. (G) Merged image showing MIOX (red) and Lipase (green) expression. White arrows indicate double-positive cells. GALK is shown in cyan and nuclei in blue. Scale bars: 20µm.

### Immune response of abdominal body wall-associated tissues to bacterial challenge

The transcriptional response of the abdominal body wall to bacterial infection was explored by challenging adult females with a systemic injection of a mixture of a Gram-negative (*E. coli)* and a Gram-positive bacteria (*M. luteus)*, or a sterile PBS control, and harvesting the abdominal body wall 4 hours later. Bulk RNA-seq analysis revealed 386 upregulated and 232 downregulated genes (logFC > 1, FDR < 0.05) in response to bacterial challenge. We observed a major induction of several antimicrobial peptides such as defensin 1 (DEF1) and cecropins (CEC1, CecB), as well as multiple components of the Toll and IMD pathways, including the transcription factor REL2 and Peptidoglycan recognition proteins (PGRPLB, PGRPLA, PGRPS1) (Supplementary Table 5). Bacterial challenge induced expression of many other immunoregulatory genes such as C-type lysozymes (LYSC1, LYSC2, LYSC7), clip-domain serine proteases (CLIPA1, CLIPA2, CLIPA4, CLIPA5, CLIPA12, CLIPA14, CLIPB5, CLIPB6, CLIPB14, CLIPB15, CLIPB17, CLIPB20, CLIPB36, CLIPC7, CLIPE1), *Anopheles Plasmodium*-responsive leucine-rich repeat proteins (APL1A, APL1B, APL1C), Leucine-rich repeat immune proteins (LRIM1, LRIM4, LRIM20), and thioester-containing proteins (TEP1, TEP2, TEP3, TEP4, TEP6, TEP15)(Supplementary Table 5). Most downregulated genes are involved in carbohydrate, amino acid, and fatty acid metabolism, as well as membrane transport (Supplementary Table 5).

To identify cell-specific responses to bacterial challenge, we compared the genes with enhanced expression after bacterial challenge in the bulk RNA-seq data with markers highly expressed in each of the cell clusters identified by snRNA-seq. Interestingly, we observed that 44 gene markers of cluster T4 were upregulated in the bulk RNA-seq data after bacterial challenge (Figure 5A), suggesting that T4 trophocytes mount a strong immune response to infection. To understand the responses of fat body and tissue-associated cells to infection, we performed ISH using probes for DEF1 and TEP1 (Figure 5B-F). TEP1 is a marker of cluster T4 and is only expressed in a subset of trophocytes in the PBS control (Figure 5B, left). However, TEP1 is expressed in all trophocytes following a bacterial challenge (Figure 5B, right). DEF1 is a shared marker of the trophocyte T4 and hemocyte (Hm1 & Hm2) clusters. We were unable to detect DEF1 expression in the control samples (Figure 5C, left). However, after bacterial challenge, DEF1 is highly expressed in a subset of trophocytes (Figure 5C, right), as well as in hemocytes (Figure 5D, lower panel), epidermal cells (Figure 5E, lower panel), and pericardial cells (Figure 5F, lower panel).

**Figure 5:**
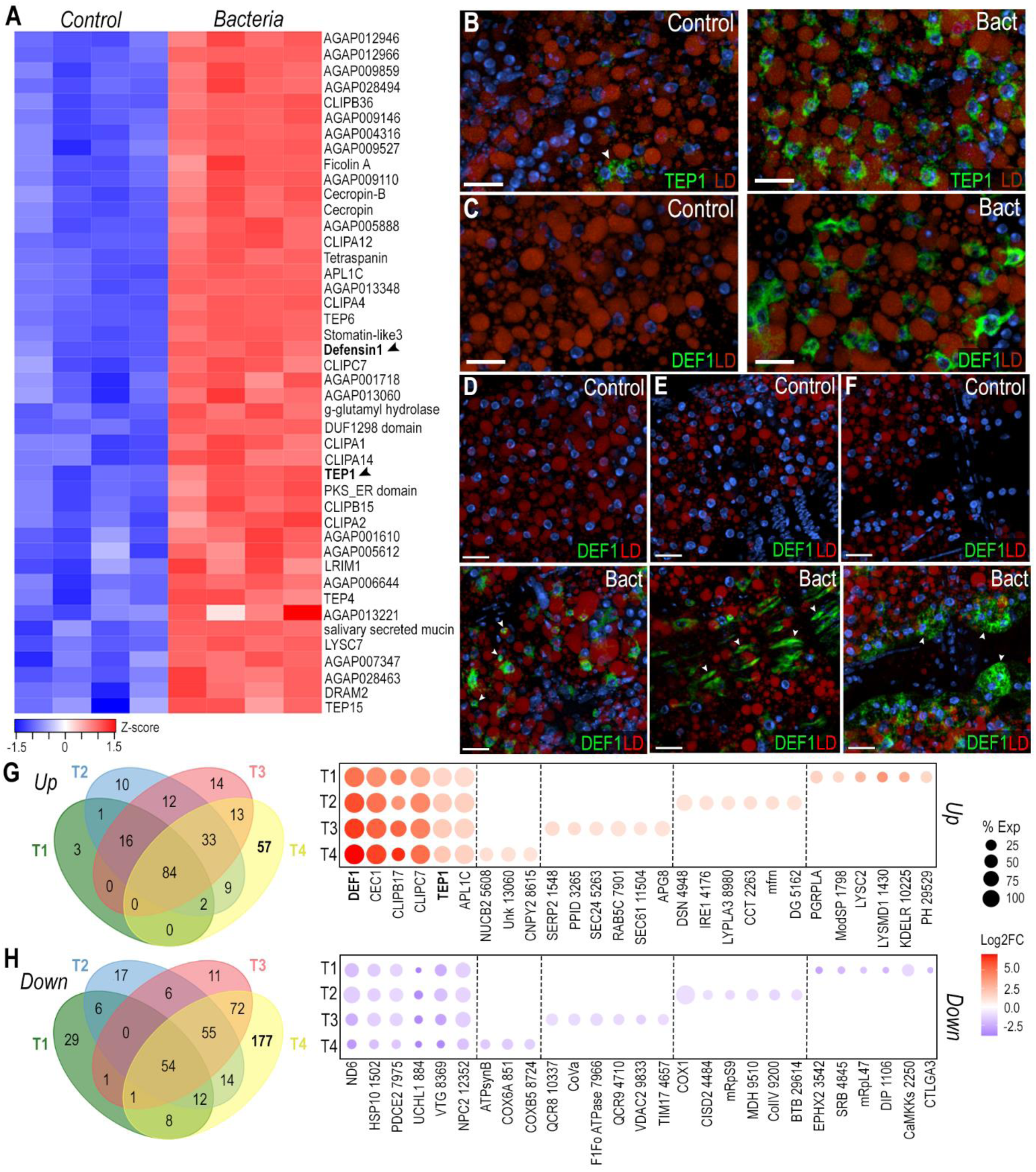
The T4 trophocyte subpopulation is activated following systemic bacterial infection. (A) Heatmap of differentially expressed genes from bulk RNA-seq analysis that are also markers of the T4 cluster. Black arrows indicate Defensin 1 (DEF1; AGAP011294) and TEP1 (AGAP010815). (B, C) RNA *in situ* hybridization of TEP1 (B) and DEF1 (C) in trophocytes under control (left) and systemic bacterial infection (right) conditions. (D - F) RNA *in situ* hybridization of DEF1 in hemocytes (D), epidermal cells (E), and pericardial cells (F) under control (top) and systemic bacterial infection (bottom) conditions. TEP1 and DEF1 are shown in green, lipid droplet (LD) in red, and nuclei in blue. White arrows indicate the corresponding cell types. Scale bars: 20µm. (G, H) Venn diagram and dot plot showing upregulated (G) and downregulated (H) genes in trophocyte subpopulations T1–T4. Dot size indicates the percentage of cells within each cluster expressing the gene, while dot color represents the log_2_ fold change. Gene names were abbreviated, with numerical identifiers corresponding to VectorBase AGAP accession numbers.

Differential expression analysis on trophocyte subpopulations revealed a core immune response with 84 genes being upregulated and 54 genes being downregulated in all trophocyte clusters (T1-T4) (Avg log2FC > 1, P val adj < 0.001) (Figure 5G-H, Supplementary Table 6). These genes with broad and strong induction included antimicrobial peptides (DEF1, CEC1, CecB), genes involved in melanization (CLIPB17, CLIPC7, CLIPB15), the complement system (TEP1, APL1C, CLIPA2), and other canonical immune genes, while mitochondrial (ND6, AGAP001502, AGAP007975) and lipid metabolism genes (AGAP000844, AGAP008369, AGAP012352) were downregulated (Supplementary Table 6). Interestingly, each trophocyte subpopulation also had specific transcriptional responses to bacterial infection (Figure 5G-H). As expected, T4 trophocytes had the strongest transcriptional response to infection, inducing expression of 57 genes, many of which are involved in immunity, such as pathogen recognition proteins (GNBPB1, PGRPLA), activation of the Toll pathway (AGAP001798), bacterial cell wall lysis (LYSC2, AGAP001430), microtubule assembly and cytoskeleton reorganization (AGAP012991, AGAP010770, Arf2), protein folding, maturation, and translocation (AGAP010225, AGAP002816, AGAP007361), and intracellular signaling (AGAP029529, AGAP010350, AGAP005352) (Figure 5G, Supplementary Table 6). Downregulated genes in cluster T4 are involved in cholesterol metabolism (AGAP003542, AGAP004845), mitochondrial translation (AGAP009737, mRpL47), transcription (AGAP001106, AGAP029767), and intracellular signaling (AGAP002250, AGAP001110) (Figure 5H, Supplementary Table 6). Most genes in T1 trophocytes were downregulated (29 genes), and 48% of them are involved in oxidative phosphorylation (Figure 5H, Supplementary Table 6). T2 trophocytes induced expression of 10 genes, 7 of which are related to protein folding, protein and vesicle transport, and autophagy (APG8) (Supplementary Table 6). Finally, T3 trophocytes increased the expression of 14 genes, some of which are involved in immunity, ER stress, or membrane remodeling (Supplementary Table 6). Differential expression analysis of hemocyte, epidermal cells, and pericardial cell clusters revealed upregulation of several core immune response genes that are also induced in the trophocyte clusters (Supplementary Table 6).

### Response of the fat body trophocytes to blood feeding

The cellular response of trophocytes to blood feeding was investigated by comparing bulk RNA-seq and snRNA-seq of the abdominal fat body between sugar-fed and blood-fed females 24 hours post-feeding (PF). In the sugar-fed sample, cluster T1 corresponds to 60.6% of the cells, while in the blood-fed sample (24 hours PF), the same cluster represents only 5.1% of the cells. Conversely, cluster T5 represented 0.1% of cells in the sugar-fed sample and becomes 54% of the cells 24 hours PF (Figure 6 A-B, Supplementary Table 2). These dramatic changes in the numbers of cells in these clusters suggest that cluster T1 undergoes such a drastic transcriptional change in response to blood feeding that the integration algorithm classifies it as a new cluster (T5), which is only present in blood-fed females. In agreement with this hypothesis, cluster T5 is greatly reduced and cluster T1 becomes predominant again 6 days PF (Figure 6C, Supplementary Table 2), when the gonadotrophic cycle has been completed, indicating that the transition of the T1 cluster to T5 is due to a prominent but transient transcriptional response to blood feeding.

**Figure 6:**
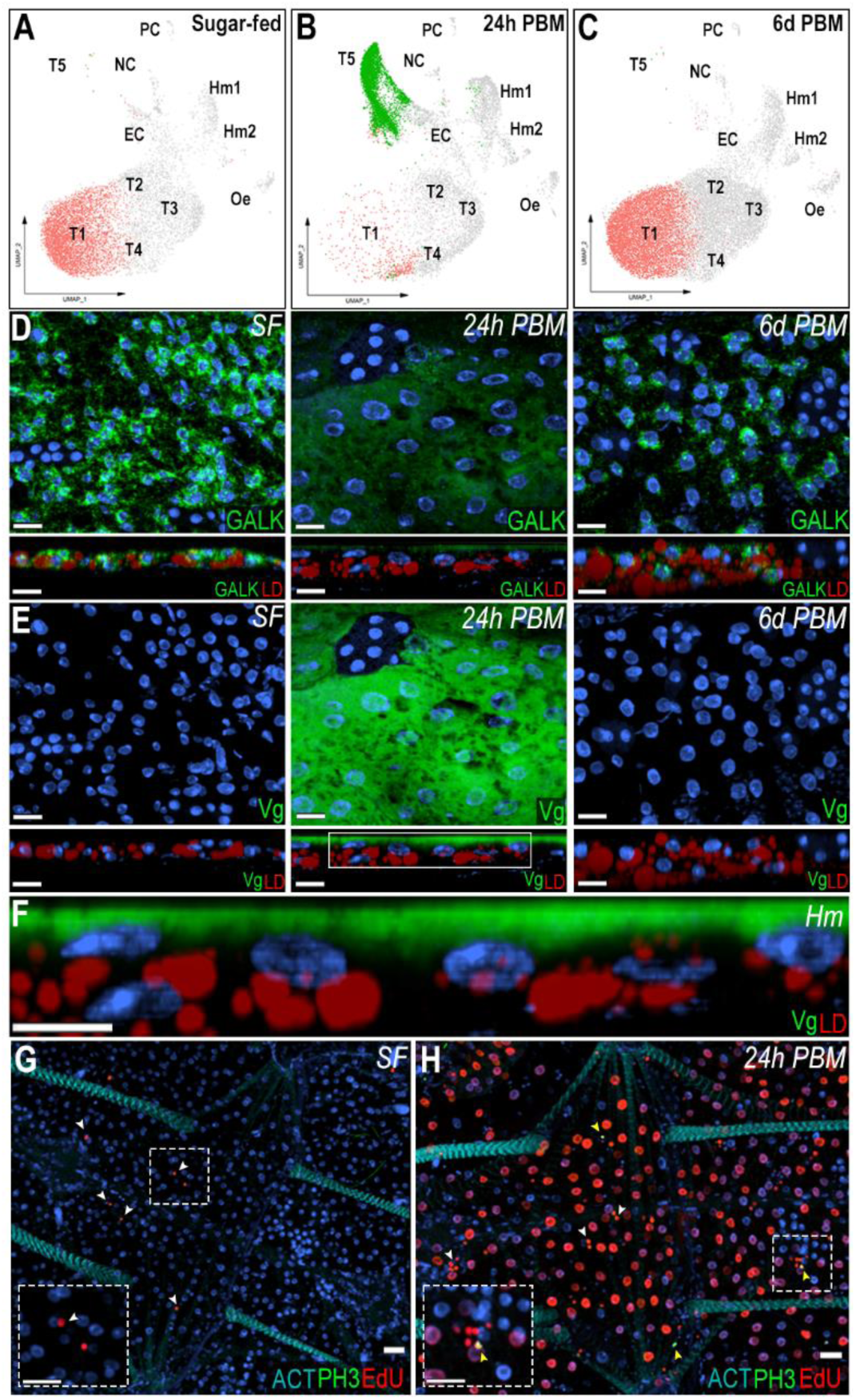
Trophocytes response to blood feeding involves intense transcriptional changes. (A-C) UMAP visualization of T1 and T5 trophocyte clusters in sugar-fed (A), 24h post-blood meal (B), and 6 days post-blood meal (C) conditions. (D, E, top) RNA *in situ* hybridization of Galactokinase (GALK; AGAP002914) (D) and Vitellogenin (Vg; AGAP004203) (E) in trophocytes on the lateral abdominal body wall under sugar-fed (left), 24h post-blood meal (middle), and 6 days post-blood meal (right) conditions. (D, E; bottom) Side view of GALK and Vg staining. Scale bars: 15µm. (F) Zoom-in of the selected region from Vg panel at 24h post blood meal. Scale bars: 10µm. GALK and Vg are shown in green, lipid droplet (LD) in red, and nuclei in blue. Hm indicate the hemolymph side.(G, H) EdU and phosphor-histone H3 (pH3) staining in sugar-fed (G) and 24 post blood meal (H) conditions. EdU is shown in red, pH3 in green, and nuclei in blue. White arrows indicate EdU-positive hemocytes and yellow arrows indicated pH3-positive hemocytes. Dashed squares show a zoom-in of hemocytes. Scale bars: 20µm.

To investigate the transition of the T1 cluster to the T5 transcriptional profile, ISH of the abdominal fat body of sugar-fed and blood-fed females collected 24 hours and 6 days post-feeding was performed. GALK was used as a general marker of sugar-fed trophocytes, with an enriched expression in the T1 cluster and vitellogenin (Vg – AGAP004203) as a marker of the T5 cluster. In sugar-fed females, GALK is expressed in the cytoplasm of all trophocytes, particularly in the perinuclear region, while Vg expression could not be detected (Figure 6D-E). In contrast, high levels of Vg expression were detected 24 hours PF, while GALK expression levels were low (Figure 6D-E). Vg is expressed at higher levels in the fat body associated with the lateral cuticle, relative to the ventral and dorsal sides of the tissue (Supplementary Figure 4). Interestingly, Vg mRNA is only detected in the layer of fat body trophocytes in contact with the hemolymph (Figure 6E-F). Furthermore, at a subcellular level, Vg mRNA is concentrated on the apical surface of the cell that faces the hemolymph (Figure 6F). The Vg mRNA subcellular localization was similar in the lateral, ventral, and dorsal fat body (Supplementary Figure 4). As expected, GALK and Vg expression 6 days PF are similar to those of sugar-fed females (Figure 6D-E).

Differential gene expression analysis between clusters T1 (sugar-fed) and T5 (24 hours PF) revealed that 171 genes were upregulated and 341 downregulated (Avg log2FC > 1, P val adj < 0.05) (Supplementary Table 7). During vitellogenesis, the fat body produces yolk protein precursors that are transported to the ovaries. Vitellogenins (AGAP004203, AGAP013109), the main yolk proteins, are highly increased in cluster T5 (Supplementary Table 7). Also, proteins involved in breaking down yolk proteins to ensure nutrient availability for the developing oocyte, such as cathepsin B precursors (AGAP004531, AGAP004534), and vitellogenic carboxypeptidase-like protein (AGAP005434) were upregulated in cluster T5 (Supplementary Table 7). Lipids are also transported from the fat body to the developing oocytes, and several proteins involved in lipid metabolism and transport, including lipophorin (AGAP001826), the main hemolymph lipid transporter, were induced in cluster T5 (Supplementary Table 7). In addition, genes related to amino acid degradation, ribosomal proteins, endoplasmic reticulum (ER) stability, protein transport, and glycosylation were increased (Supplementary Table 7). Meanwhile, genes related to autophagy, carbohydrate metabolism, circadian clock, immunity, signaling, and transcription were downregulated (Supplementary Table 7).

Bulk RNA-seq analysis confirmed a strong transcriptional response to blood feeding, with 1,655 genes upregulated and 1,368 genes downregulated (logFC > 1, FDR < 0.01) (Supplementary Table 8). In agreement with the differential gene expression of cluster T5, vitellogenin (AGAP004203 had a 5,898-fold increase), cathepsin B precursor (AGAP004534, 121-fold increase), and expression of vitellogenic carboxypeptidase proteins (AGAP005434, 2.8-fold increase; AGAP007505, 2.8-fold increase) was also increased 24 h PF (Supplementary Table 8). Interestingly, increased expression of genes involved in DNA replication, such as DNA replication licensing factors (MCM3, MCM4, MCM5, MCM6), DNA polymerase subunits (10 different subunits), replication factors (A1, A2, A3, and 3 C-subunits), proliferating cell nuclear antigen (PCNA), ribonuclease H2 subunit B, helicase Dna2, and DNA ligase 1 was observed; as well as several genes related to DNA repair (MSH2, Rad62, RAD50, MRE11, ERCC1), and cell cycle such as cyclins (A, C, H, T) and cyclin-dependent kinases (CDK1, CDK4) (Supplementary Table 8).

To determine which cells associated with the abdominal wall were undergoing DNA replication in response to blood feeding, 5-ethynyl-2’-deoxyuridine (EdU) was injected into the hemolymph of sugar-fed or blood-fed females, 6 hours PF. Samples of sugar-fed females and blood-fed females were analyzed 18 hours after EdU injection. We were not able to detect EdU incorporation in trophocytes of sugar-fed females (Figure 6G). In contrast, the majority of trophocytes from blood-fed females incorporated EdU in their DNA, resulting in strong nuclear staining (Figure 6H). A few hemocytes were also EdU^+^ in both sugar-fed and blood-fed mosquitoes (Figure 6G-H, white arrows). EdU staining was not detected in any other cell types. Staining for phosphohistone H3 (pH3), a marker of mitosis, was negative in trophocytes of sugar-fed and blood-fed females (Figure 6G-H). Only a few pH3^+^ hemocytes (Figure 6H, yellow arrows) were observed in the body wall of blood-fed mosquitoes; all other cells were negative. Blood feeding also triggered a strong downregulation of genes involved in carbohydrate metabolism, such as genes involved in glycolysis, citric acid cycle, beta oxidation, elongation of very long fatty acids, ribosomal biogenesis, mitochondrial organization, protein folding, and the TOLL pathway (Supplementary Table 8).

## Discussion

Single-cell transcriptome analysis has transformed our understanding of cellular heterogeneity by enabling the analysis of individual cells and has provided new insights into cell-specific functions, morphology, and dynamic transcriptional states. In this study, we used snRNA-seq to investigate the transcriptional landscape of the abdominal fat body and associated tissues of *An. gambiae* females to overcome the technical challenge of isolating intact single cells from a tissue with very high lipid content. The protocol involves a delipidation step, as lipids interfere with cell dissociation and 10X single-cell partitioning and Gel Bead-in-Emulsion (GEM) formation. Previous studies have demonstrated that the nuclear transcriptome correlates with the cytoplasmic mRNA composition [26, 27], making snRNA-seq a viable alternative for transcriptomic profiling when whole-cell isolation is not feasible.

Previous single-cell and single-nucleus transcriptomic studies in insects have begun to illuminate the cellular complexity of metabolically and immunologically active tissues such as the fat body. In *Drosophila melanogaster*, snRNA-seq has been used to resolve cell-type-specific responses to immune challenge and aging, revealing subpopulations of fat body cells with specialized metabolic or inflammatory functions [28]. snRNA-seq was applied to *Bombyx mori* to characterize distinct metabolic and immune programs in fat body cells during viral infection, also using nuclear-based transcriptional profiling [29]. More recently, scRNA-seq and metabolomics were used to investigate the transcriptional and metabolic responses of the *Aedes aegypti* fat body and midgut following a blood meal, uncovering functionally distinct cell types and dynamic responses to blood feeding [30]. Despite these advances, a comprehensive understanding of fat body cell heterogeneity and its response to infection and immune priming has not been available for anopheline vectors of malaria.

Our observation of a high number of hemocytes closely associated with the fat body (Figure 2C, Supplementary Table 2) underscores the importance of local immune surveillance in this tissue. We found that while granulocytes typically represent only a small fraction (1–5%) of circulating hemocytes [20, 21], they account for nearly 87% of fat body-associated hemocytes (Supplementary Figure 2, Supplementary Table 2), suggesting that a substantial proportion of granulocytes embedded in the trophocyte layers of the tissue may represent resident immune cells. This supports the growing recognition tissue-resident immune cells in insects that respond readily to local infections [31–33]. In contrast, granulocytes loosely attach on the hemolymph side of fat body basal lamina and circulating oenocytoids, which are more abundant in circulation and underrepresented in the fat body, may have a more systemic role and mobilize to remote sites of damage or infection.

We observed a doubling of the number of fat body-associated granulocytes in response to bacterial challenge, suggesting either proliferation of resident cells and/or recruitment from circulation. This aligns with previous reports of hemocyte accumulation in the periosteal regions post-infection [32] and supports the idea that immune challenge triggers dynamic shifts in hemocyte distribution and abundance. Additionally, the increased capture of both granulocytes and oenocytoids after a blood feeding agrees with previous reports of increased number of circulating hemocyte in blood-fed females, illustrating dynamic changes in tissue-associated hemocyte populations [34]. The presence of pH3- and EdU-positive hemocytes (Figure 6G-H), indicating mitotic activity and DNA synthesis within these cells is indicative of cell proliferation. Together, these findings highlight the plasticity of mosquito hemocyte populations and suggest that the fat body serves not only as an immune-responsive tissue but also as a compartment for immune cell residence and activity.

Oenocytes play a key role in the immune priming response. In *Aedes aegypti*, they represent approximately 5.9–9.8% of fat body cells in adult females [7]. We observed a strong increase in fatty acid synthase mRNA levels in oenocytes from primed females (Figure 3C-D). We confirmed that the DBLOX mRNA is expressed in oenocytes, and that modest increase in DBLOX mRNA expression previously reported in primed females (1.5-2.0-fold) is due to the increase in oenocyte numbers [18], as we did not observe increased gene expression by ISH on individual cells from primed females. Although hemocyte numbers did not change, we do observe a distinct transcriptional response characterized by increased expression of extracellular matrix and adhesion-related genes (Supplementary Table 2, Supplementary Table 4) in primed females. This suggests enhanced hemocyte patrolling or stronger interaction with the fat body tissue. Additionally, upregulation of genes involved in long-chain fatty acid synthesis and cholesterol metabolism in hemocytes suggests that they may also synthesize bioactive lipids in response to immune priming (Supplementary Table 4).

Of the five trophocyte clusters that were identified (Figure 1B), clusters T1 and T2 shared similar transcriptomes with subtle differences in gene expression (Figure 1C), indicative of a basal metabolic state. In contrast, clusters T3, T4, and T5 exhibited specialized gene expression patterns: T3 was enriched for metabolic genes, T4 for immune-related genes, and T5 for vitellogenesis-related genes (Supplementary Table 1, Figure 6A-C). Our findings reveal that the mosquito mounts a robust and multilayered immune response to bacterial infection, involving not only trophocytes but also hemocytes, epidermal cells, and pericardial cells (Figure 5B-F). The activation of both TOLL and IMD pathways, along with the induction of pattern recognition receptors, melanization components, and antimicrobial effectors, points to a broad engagement of classical innate immune mechanisms. Importantly, the identification of a core set of immune genes expressed across multiple cell types suggests a conserved, systemic response to infection. This shared transcriptional program is complemented by cell-type-specific immune signatures, indicating functional specialization that may allow for spatial or temporal modulation of the immune response. The expression of immune genes in T4 trophocytes, even in the absence of infection, raises the possibility that this subpopulation plays a role in immune surveillance or maintaining immune readiness—functions that parallel tissue-resident immune cells in vertebrates. These immune responses appear to come at a metabolic cost, as evidenced by the downregulation of oxidative phosphorylation-related genes. This trade-off between immunity and metabolism is consistent with findings in other insects and vertebrates, where immune activation often leads to mitochondrial remodeling or dysfunction, either as a host-driven adaptation or as a result of pathogen interference [35–37].

In our study, trophocytes transitioned from a basal state (cluster T1) to a highly active state (cluster T5) characterized by dramatic changes in gene expression (Figure 6A-E, Supplementary Table 7, Supplementary Table 8). Notably, this shift was reversible by six days post-feeding, suggesting a temporary transcriptional reprogramming rather than permanent differentiation. Trophocyte endoreplication is well-documented in mosquitoes and other insects, serving to increase transcriptional output in support of reproduction [12, 38–40]. Our findings reinforce this model by providing both molecular and histological evidence, as we observed EdU-positive, but pH3-negative trophocytes (Figure 6G-H), consistent with DNA replication in the absence of mitosis. This aligns with a pronounced transcriptional induction of genes involved in DNA replication and cell cycle regulation at 24 hours post-blood feeding (Supplementary Table 8). These data suggest that endoreplication is a mechanism that enables a rapid response to the extreme metabolic demands that allows trophocytes to amplify expression of yolk protein genes and other genes involved in oogenesis. More broadly, these results highlight the dynamic and reversible transcriptional plasticity of the fat body and emphasize the importance of considering endocycling states when interpreting insect single-nucleus transcriptomic data.

Ultrastructural studies of the fat body in *Aedes aegypti* reveal significant changes in trophocytes as they transition from the previtellogenic to the vitellogenic phase following blood feeding. During this transition, ribosomes that were initially free in the cytoplasm associate with the endoplasmic reticulum (ER), the Golgi complex enlarges and associates with the rough endoplasmic reticulum (RER), and the surface of the plasma membrane increases by forming more invaginations [7, 12]. Interestingly, these changes are more pronounced in trophocytes adjacent to the hemolymph, suggesting that these cells are more actively engaged in protein synthesis and secretion towards the hemolymph [7, 12], and agrees with the localized expression of Vg mRNA we observed by ISH in this trophocytes.

Furthermore, at a subcellular level, Vg transcripts are localized in the apical region of the trophocyte layer facing the hemolymph. In mammals, mRNA localization is often regulated by RNA-binding proteins associated with the cytoskeleton, influencing translation efficiency through ribosome positioning [41, 42]. Similar regulatory mechanisms may be at play in trophocytes, enabling efficient protein export into the hemolymph. Notably, higher expression of Vg is observed in the fat body associated with the lateral body wall. This spatial pattern may be influenced by the substantial mechanical stretching that the mosquito lateral body wall undergoes during blood feeding, potentially affecting local gene expression. The lateral lobes of the fat body are also more prominent [44], which may increase surface area contact with the hemolymph, thereby enhancing Vg export.

In summary, our snRNA-seq analysis reveals the remarkable transcriptional complexity and functional specialization of the mosquito fat body in response to blood feeding and infection. By overcoming technical barriers to nucleus isolation, we uncovered a dynamic interplay between metabolic and immune pathways across multiple fat body-associated cell types. Hemocytes appear to be locally enriched and transcriptionally responsive, suggesting a model of tissue-resident immune surveillance. Trophocytes exhibit distinct transcriptional states, with a reversible transition toward reproductive specialization and endoreplication in response to blood feeding. Notably, the polarized expression of vitellogenin mRNA and metabolic shifts underscore the spatial and functional plasticity of fat body trophocytes. Together, these findings support a model in which the fat body integrates metabolic, reproductive, and immune functions through spatially organized and temporally regulated transcriptional programs. This study provides a high-resolution atlas of the mosquito fat body and associated cells, revealing key physiological responses to blood feeding, bacterial challenge, and immune priming, which establish a foundation for future efforts to manipulate vector competence by targeting tissue-specific or state-dependent gene expression.

## Methods

### Ethics statement

Public Health Service Animal Welfare Assurance #A4149-01 guidelines were followed according to the National Institutes of Health Animal (NIH) Office of Animal Care and Use (OACU). These studies were done according to the NIH animal study protocol (ASP) approved by the NIH Animal Care and User Committee (ACUC), with approval ID ASP-LMVR5.

### Mosquito rearing

*Anopheles gambiae* mosquitoes (G3 strain – CDC) were maintained under standard insectary conditions at 28°C, 80% relative humidity, and a 12:12 hour light:dark cycle. Adult stages were fed *ad libitum* with cotton pads soaked in 10% Karo syrup. Eggs were hatched in plastic trays containing 1L of deionized water supplemented with a pinch of powdered koi fish food and 10mL of yeast solution (prepared by dissolving two tads of live baker’s yeast in 50mL of deionized water). Larvae stages were reared in trays with 1L of deionized water and fed every other day with fish food pellets (Tetramin, Blacksburg, VA, USA). The colony was maintained by feeding adult female mosquitoes on defibrinated cow blood using an artificial membrane feeding system.

### Blood feeding

Female *An. gambiae* mosquitoes, 4-5 days post-eclosion, were blood-fed on 3- to 4-week-old female BALB/c uninfected mice (Charles River, Wilmington, MA, USA) for 15 minutes. Prior to feeding, mosquitoes were starved for approximately 16 hours. Only fully engorged females were selected and maintained on 10% karo syrup following feeding until sample harvesting. Same-age female mosquitoes maintained on *ad libitum* 10% karo syrup were used as a sugar-fed control. Abdominal fat body tissues were dissected 24 hours post-blood meal, along with their age-matched sugar-fed control mosquitoes. Mice were housed in individually ventilated cages at 20–21°C and 56% relative humidity with a 12:12 hour light-dark cycle and provided with sterile water and food *ad libitum*.

### *Plasmodium berghei* infection

Mosquito infections with *Plasmodium berghei* were carried out using the transgenic GFP-expressing ANKA GFPcon 259cl2 parasite line [45], maintained by serial passages into 3- to 4-week-old female BALB/c mice from frozen stocks. Prior to mosquito feeding, parasitemia was monitored using *in vitro* exflagellation assays as previously described [46]. Four-to-five-day-old female mosquitoes were allowed to feed on infected mice once parasitemia reached 3-6% and 1-2 exflagellations per field were observed. Same-age uninfected mice were used to feed control group of blood-fed mosquitoes. After feeding, both control and infected mosquitoes were maintained at 19°C, 80% humidity, and 12:12 hour light:dark cycle. After 48h, mosquitoes were shifted to 28°C, under the same humidity and light conditions, to reduce oocyst development. Female mosquitoes were allowed to lay eggs at 4 days post-blood meal. Abdominal fat body tissues from control and infected samples were dissected 6 days post-blood meal.

### Bacteria challenge

*Escherichia coli* and *Micrococcus luteus* were kept as frozen stocks at −80°C. A pre-inoculum for each bacterium was set up and allowed to grow overnight at 37°C, 250 rpm in a shaker incubator. On the day of the experiment, the pre-inoculum was diluted in fresh LB media and allowed to grow in the same conditions until OD_600_ of 0.5. *E. coli* and *M. luteus* cultures were mixed at a 1:1 ratio and washed with sterile PBS to remove toxins. Adult 4-day-old female mosquitoes were injected with 138 nL of sterile PBS or bacteria mix. Abdominal fat body tissues from control and infected samples were dissected 4 hours post-injection.

### Single nuclei isolation

Female mosquitoes were cold-anesthetized on ice and placed on a drop of filter-sterilized adult hemolymph-like saline (HLS) buffer (Calcium chloride 2mM, Potassium chloride 5mM, HEPES pH7.4 5mM, Magnesium chloride 8.2mM, Sodium chloride 108mM, Sodium bicarbonate 4mM, Sodium phosphate 1mM, sucrose 10mM, Trehalose 5mM – [47]). To isolate the abdominal fat body tissue, the abdomens were separated from the thorax. The gut, ovaries, and Malpighian tubules were then removed from the body cavity, leaving the abdominal fat body attached to the cuticle. For each experimental condition, two pools of 25 tissues were dissected and gently squeezed on ice in HLS buffer with a plastic pestle. The samples were washed twice with 80% methanol to fix and remove lipids. Centrifugations were carried out for 5 min at 500 x g at 4°C. To release the nuclei, 500µL of lysis buffer (1mL Nuclei PURE lysis buffer, 1ul 1M DTT, and 10uL Triton X-100; Sigma, MO, USA) was added to each sample and homogenized using a pre-cooled 1mL Dounce tissue grinder, applying 15 strokes with both the loose and tight pestle. Samples were gently mixed with 1.8mL of a 1.8M sucrose cushion solution (5.4mL Nuclei PURE 2M Sucrose Cushion solution and 0.6mL Nuclei PURE Sucrose Cushion Solution; Sigma, MO, USA) to dilute the lysis buffer. The sample was then subjected to a one-step continuous sucrose cushion for further purification. Each sample was divided into two tubes, with 900µL of the homogenized sample carefully layered over 500µL of 1.8M sucrose cushion solution. Blood-fed samples were divided into four tubes. Centrifugation was carried out at 13,000 x g for 45 minutes at 4°C. The supernatant was gently discarded, and the resulting pellet, containing the nuclei, was washed twice with washing buffer (1% BSA in PBS + 0.2u Protector RNase Inhibitor; Sigma, MO, USA). All solutions were kept on ice during the procedure. Finally, the nuclear suspension was stained with 0.4% trypan blue, and the nuclear yield was estimated using a hemocytometer. The targeted concentration of nuclear suspension was 700-1200 nuclei per microliter.

### Chromium 10X snRNA-seq library preparation and sequencing

After performing the single nuclei isolation of the fat body of *An. gambiae* female mosquitoes with different stimuli, an estimated number of 10,000 fresh single nuclei were immediately used for 10X Genomics Chromium droplet snRNA-seq. Single nuclei encapsulation, cDNA synthesis, and libraries preparation were carried out using the Chromium Next GEM Single Cell 3ʹ Reagent Kits v3.1 (10x Genomics, Pleasanton, CA), following the manufacturer’s instructions. Assessment of cDNA and library quantification and quality was performed using 4200 Agilent TapeStation System with a High Sensitivity D5000 ScreenTape (Agilent, Santa Clara, CA). Paired-end sequencing was performed with Illumina NextSeq 550 or NextSeq 2000 platforms.

### Single-Nucleus RNA-seq Processing and Ambient RNA Correction

To generate the single-nuclei gene expression matrix, raw sequencing reads were aligned to the *Anopheles gambiea* PEST (AgamP4.14, v59) reference genome using the Cell Ranger pipeline (version 7.0.0, 10× Genomics, Pleasanton, CA). The reference genome and corresponding annotation file were obtained from VectorBase (https://vectorbase.org/common/downloads/Current_Release/AgambiaePEST). The gene annotation (gtf) file was filtered to retain protein-coding genes, and a custom reference was built using the “cellranger mkref” function. Alignment and quantification were performed with the “cellranger count” function, using the parameters “--include-introns=true” and “--expect-cells=10000”, to generate both raw and filtered gene-barcode matrices. As nuclei isolation can result in the presence of ambient RNA—potentially from lysed cells—that becomes incorporated into droplets containing intact nuclei, we applied ambient RNA correction to the dataset. Correction was performed using the decontX algorithm from the celda package [48], which models and removes contamination from ambient background RNA in snRNA-seq data.

### Quality control and cell filtering

Quality control and cell filtering were performed using the Seurat R package (version 4.2.1). For each sample, nuclei were retained based on sample-specific thresholds for the number of detected genes (Features), the percentage of mitochondrial gene expression, and the percentage of ribosomal gene expression, to remove low-quality nuclei and potential doublets. Cells were included if they expressed between 200 and 2000 genes for most samples (Naïve 1, Naïve 2, Prime 1, Prime 2, and Blood-fed), with adjusted upper bounds for PBS-control, bacteria-injected (2500), and Sugar-fed samples (1500). To minimize the inclusion of damaged or stressed nuclei, cells with a mitochondria gene percentage exceeding 2.5% (Naïve 1 and Prime 1), 1.5% (Naïve 2 and Prime 2), 7% (PBS-control), 5% (Bacteria-injected), or 10% (Sugar-fed and Blood-fed) were excluded. Additionally, a ribosomal gene expression threshold was applied, with upper limits ranging from 10% to 20% across samples to account for potential variation in ribosomal content between conditions. Cellular complexity, assessed using the log10GenesPerUMI metric, was also evaluated to ensure transcriptomic diversity and minimize amplification bias; all retained nuclei had a complexity score greater than 0.75. These filtering parameters were chosen to optimize the retention of high-quality nuclei while minimizing technical artifacts and ambient RNA contamination.

### Integrative analysis, cell clustering, and marker gene identification

Seurat-based integration and clustering analysis was performed on previously described datasets: Naïve and Prime, PBS-control and bacteria-injected, and Sugar-fed and Blood-fed. This was achieved by first log-normalizing the gene counts of the remaining filtered nucleus using the “NormalizeData” function. Next, the top 2000 most variable genes were identified for each sample using the “FindVariableFeatures” function and tested for suitability as integration anchors using the “FindIntegrationAnchors” function. A final set of 2000 anchors were then used in the “IntegrateData” function to integrate the samples into a single dataset. Following integration, the dataset was scaled using the “ScaleData” function. Dimensionality reduction was performed using “RunPCA”, and the first 30 principal components were used to generate a two-dimensional embedding with “RunUMAP”. Clustering was conducted with “FindNeighbors” and “FindClusters”, applying a resolution 0.3 to identify transcriptionally distinct cell populations. Marker genes for each cluster were identified using “FindAllMarkers” function in Seurat, requiring that genes must be detected in at least 25% of cells of the integrated dataset, an average log2 fold change (Avg log2FC) ≥ 0.5 and, and an adjusted p-value (P val adj) < 0.05. Differential gene expression analysis was performed using the “FindMarkers” function to assess transcriptomic changes between treatment groups. For comparisons between PBS-control and bacteria-injected, as well as Sugar-fed and Blood-fed samples, stringent thresholds were applied, an average log2 fold change (Avg log2FC) ≥ 1 or ≤ −1 and a p adjusted value (P val adj) < 0.001. A more permissive threshold, Avg log2FC ≥ 0.5 and P val adj < 0.05, was used for comparisons between *P. berghei*-infected and naïve samples. Raw sequencing data have been deposited in the SRA database under BioProject accession number PRJNA1288431 (https://dataview.ncbi.nlm.nih.gov/object/PRJNA1288431?reviewer=he84ttfsjqcmttfp3ccg035vi8).

### Bulk RNA-seq RNA extraction, library preparation, and sequencing

Body wall tissues (abdominal fat body attached to the cuticle) were dissected from adult female mosquitoes as previously described. Each biological replicate consisted of a pool of 10 tissues (n =4). Dissected tissues were immediately transferred to 300µL of TRIzol^TM^ reagent (ThermoFisher Scientific, MA, USA) and stored at −80◦C until processing. Tissues were homogenized using an electric homogenizer, followed by the addition of 700µL of TRIzol^TM^ reagent. Samples were centrifuged at 12,000 x g for 5 minutes at 4°C to pellet the cuticle. After a 5-minute incubation at room temperature, 200uL of chloroform was added, and samples were vortexed briefly. Following a 3-minute room temperature incubation, samples were centrifuged at 12,000 x g for 15 minutes at 4°C. The aqueous phase was transferred to a fresh 1.5mL microcentrifuge tube, and the RNA was precipitated by adding 500µL of isopropanol. Samples were gently mixed by inversion, incubated on ice for 15 minutes, and centrifuged at 12,000 x g for 15 minutes at 4°C. The resulting pellet was washed with 1mL of 75% ice-cold ethanol, briefly vortexed, and centrifuged at 7,500 x g for 5 minutes at 4°C. Supernatant was discarded, and pellets were air-dried before resuspension in 30µL of nuclease-free water. RNA was homogenized and incubated at 55°C for 5 minutes to enhance solubilization. Final RNA samples were stored at −80°C. RNA libraries were prepared with NEB Next Ultra II RNA Library prep kit (New England Biolabs), and sequencing was performed on Illumina NovaSeq 6000 sequencer (150bp paired-end reads) by Novogene (Sacramento, CA). Raw sequencing data have been deposited in the SRA database under BioProject accession number PRJNA1288431 (https://dataview.ncbi.nlm.nih.gov/object/PRJNA1288431?reviewer=he84ttfsjqcmttfp3ccg035vi8).

### Bulk RNA-seq data and statistical analysis

The quality of raw Illumina reads was checked using the FastQC tool (version 0.12.0) (https://www.bioinformatics.babraham.ac.uk/projects/fastqc/). Low-quality sequences with a Phred quality score (Q) below 20 and the Illumina adaptors were removed using TrimGalore v.0.6.7 (https://github.com/FelixKrueger/TrimGalore). High-quality reads were aligned to the *Anopheles gambiea* PEST (AgamP4.14, v59) transcriptome using RSEM v.1.3.3 (Li and Dewey 2011) with Bowtie 2 as the aligner for transcript quantification. The reference transcriptome was obtained from VectorBase (https://vectorbase.org/common/downloads/Current_Release/AgambiaePEST). Downstream analysis, including multidimensional scaling (MDS) plots and pairwise differential expression analysis were carried out with the edgeR package (Robinson, McCarthy and Smyth 2010) for R. For the differential expression analysis, the raw read counts of each transcript were used as input for edgeR, followed the standard TMM-normalization. Transcripts were considered differentially expressed if they exhibited an absolute LogFoldChange (logFC) ≥ 1 and a false discovery rate (FDR) < 0.05. For comparisons involving naïve and *P. berghei*-challenged samples, a relaxed threshold of logFC ≥ 0.5 with FDR < 0.05 was applied.

### In situ hybridization (ISH)

Female mosquitoes were cold-anesthetized and placed on a drop of PBS. The abdomens were detached from the thorax and the midgut, ovaries, and Malpighian tubules were carefully removed from the body cavity. The body wall, including the abdominal fat body attached to the cuticle, was then incised along the lateral side using fine scissors. The dissected tissues were fixed for 1 hour in 4% paraformaldehyde on a 9-well plate at room temperature. After fixation, the tissues were gently transferred to a 1.5mL microcentrifuge tube and subjected to a series of washes. Tissues were washed three times for 5 minutes each with 1mL of PBST (0.1% Triton X-100 in PBS), with gentle rocking at room temperature. To reduce lipid content, the tissues were washed with increasing concentrations of methanol (PBST: Methanol; 7:3, 1:1, 3:7) for 5 minutes at room temperature. An absolute methanol wash was then performed for 10 minutes, after which the tissues were rehydrated by washing with decreasing concentrations of methanol (PBST: Methanol; 3:7, 1:1, 7:3) for 5 minutes each. Finally, tissues were washed four times with PBST for 5 minutes each. ISH was performed using the RNAscope Multiplex Fluorescent Reagent Kit v2 assay (ACDBio, Abingdon, United Kingdom), adapted from the protocol described by [21]. Tissues were permeabilized with Protease Plus, provided by the kit, for 30 minutes at room temperature. Following two washes with PBST for 5 minutes, hybridization was carried out for 2 hours at 40°C. Amplification of the hybridized probes signal step was performed in sequential steps: AMP1 and AMP2 for 30 minutes each and AMP3 for 15 minutes, all at 40°C. Between each amplification step, tissues were washed twice with wash buffer for 2 minutes at room temperature. Fluorescent signal development was achieved using the TSA-based Opal 4-Color Automation IHC kit (PerkinElmer, MA, USA). RNA probes specific to the target genes were synthesized by ACDBio (Abingdon, United Kingdom). Fluorescent labeling was caried out stepwise, with the addition of corresponding HRP (C1, C2, C3, or C4) for 15 minutes at 40°C. Following each HRP incubation, tissues were washed twice for 2 minutes before incubation with the Opal fluorophore for 30 minutes at 40°C. After each fluorophore incubation, tissues were washed twice for 2 minutes, and HRP activity was blocked by a 15-minute incubation with HRP blocker at 40°C. For morphological visualization, tissues were stained with Hoechst (1:10,000) to label the nuclei, and BODIPY 488nm (1:5,000 – ThermoFisher Scientific, MA, USA) to visualize neutral lipids in wash buffer for 10 minutes. Following a final wash to remove excess Hoechst and BODIPY, slides were mounted in a drop of ProLongTM Gold Antifade Mountant (ThermoFisher Scientific, MA, USA). Tissues were gently positioned and flattened in Prolong under a dissecting microscope to avoid folding or overlapping, and slides were allowed to dry overnight at room temperature in the dark.

### Confocal microscopy and image processing

Confocal images were captured using a Leica TCS SP8 (DM8000) confocal microscope (Leica Microsystems, Wetzlar, Germany) with either a 20X or a 40X oil immersion objective equipped with a photomultiplier tube/ hybrid detector. Abdominal fat bodies were visualized with a white light laser, using 488 nm excitation and 52nm emission for BODIPY 488 dye (neutral lipids), 550 nm excitation and 570nm emission for Opal 570 (ISH probes), 588 nm excitation and 620nm emission for Opal 620 (ISH probes), 676 nm excitation and 690nm emission for Opal 690 (ISH probes), and a 405 nm diode laser for nuclei staining (Hoechst 33342). Images were taken using sequential mode and variable z-steps. Image processing and merging were performed using Imaris 10.0.0 (Bitplane, MA, USA) and Adobe Photoshop CC (Adobe Systems, CA, USA).

## Data Availability

All data that support this study are available from the corresponding authors upon request. Raw fastq files from single nucleus and bulk RNA sequencing are available the SRA database under the Bioproject number PRJNA1288431. All images used in this manuscript are available from the corresponding authors upon request.

## Acknowledgments

We thank Kevin Lee, Yonas Gebremicale, and André Laughinghouse for insectary support. We thank Nathania Dabila and Mu Jianbing for sequencing support. This work was funded by the Intramural Research Program of the Division of Intramural Research Z01AI000947, NIAID.

## Authors Contributions

Conceptualization: SSDC, CB-M, Methodology: SSDC, ABFB, CM, Investigation: SSDC, ABFB, CM, Visualization: SSDC, ABFB, Funding acquisition: CB-M, Project administration: CB-M, Supervision: CB-M, Writing—original draft: SSDC, CB-M, Writing—review & editing: SSDC, CB-M.

