## Supplementary figures for "Cell-specific responses of *Anopheles gambiae* fat body to blood feeding and infection at a single nuclei resolution"

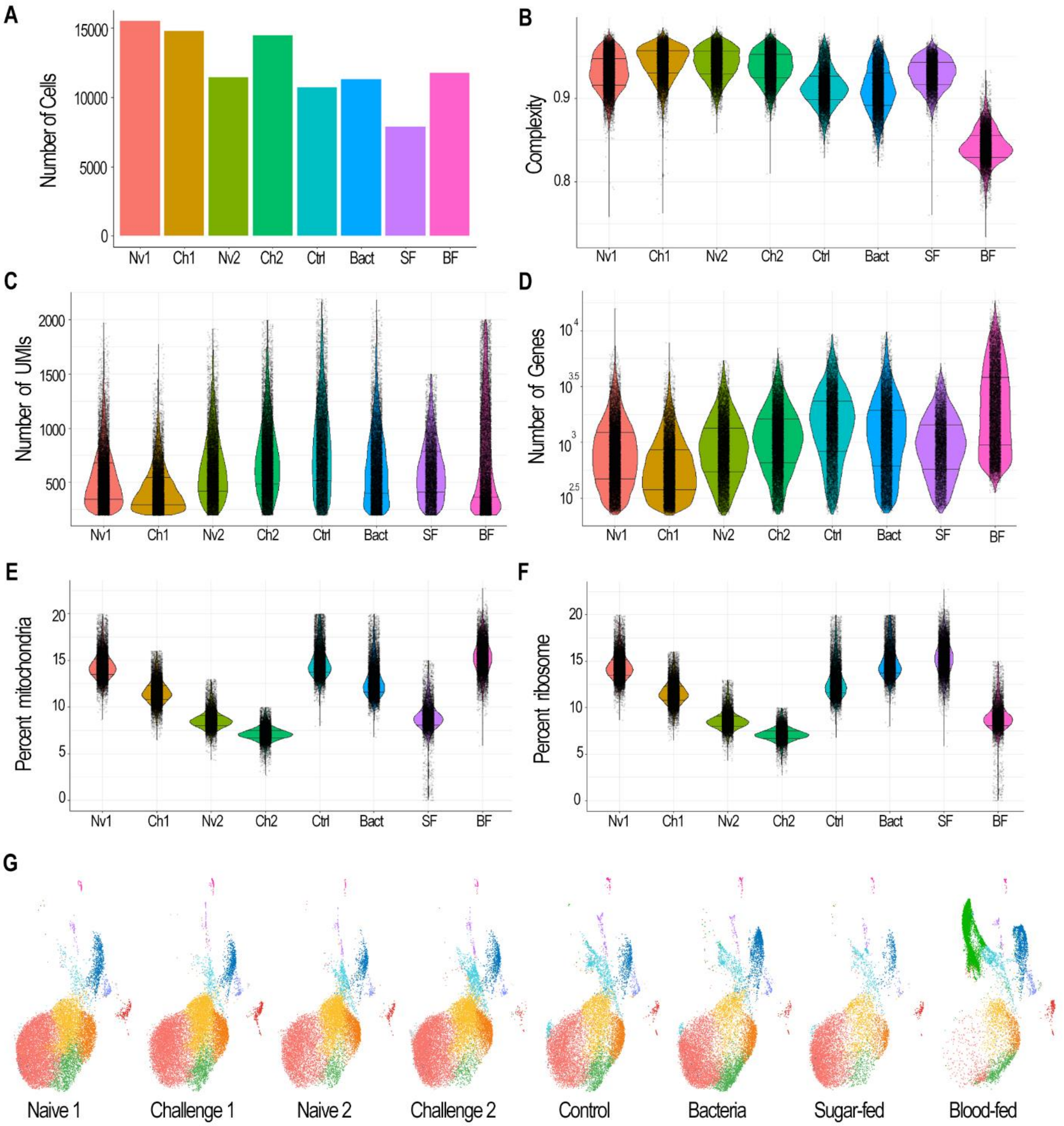


**Supplementary Figure 1. Quality control metrics and clustering analysis of *Anopheles gambiae* abdominal body wall single-nucleus RNA-seq data.** (A) Total number of cells captured per sample. (B) Distribution of complexity per sample, represented as log10GenesPerUMI. (C) Total number of UMIs per sample. (D) Number of detected genes (features) per sample. (E) Percentage of mitochondrial gene expression per sample. (F) Percentage of ribosomal gene expression per sample. (G) UMAP visualization of integrated abdominal body wall nuclei, shown separately for each sample, at clustering resolution 0.3.

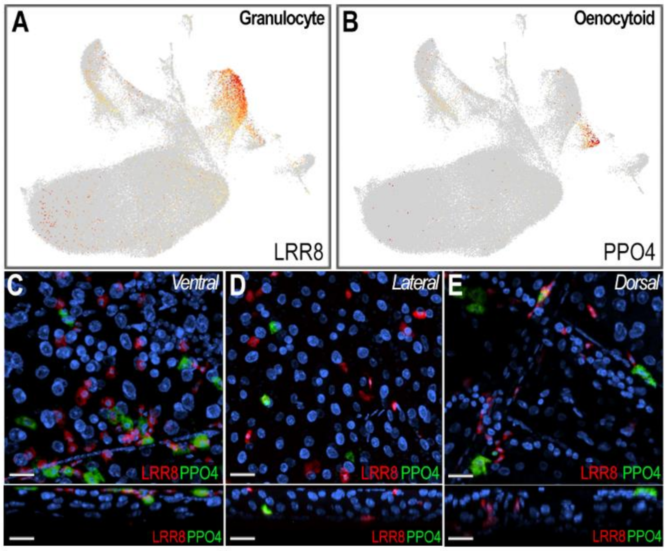


**Supplementary Figure 2. Localization and characterization of fat body-associated hemocytes.** (A) Feature plot showing expression of Leucine-rich repeat 8 (LRR8; AGAP004017), a marker of the granulocytes (cluster Hm1). (B) Feature plot showing expression of prophenoloxidase 4 (PPO4; AGAP004981), a marker of the oenocytoids (cluster Hm2). (C-E) RNA *in situ* hybridization of LRR8 and PPO4 on ventral (C), lateral (D), and dorsal (E) regions of abdominal fat body tissue. (C-E, bottom) Corresponding side views of each region. LRR8 is shown in red, PPO4 in green, and nuclei in blue. Scale bar: 20µm.

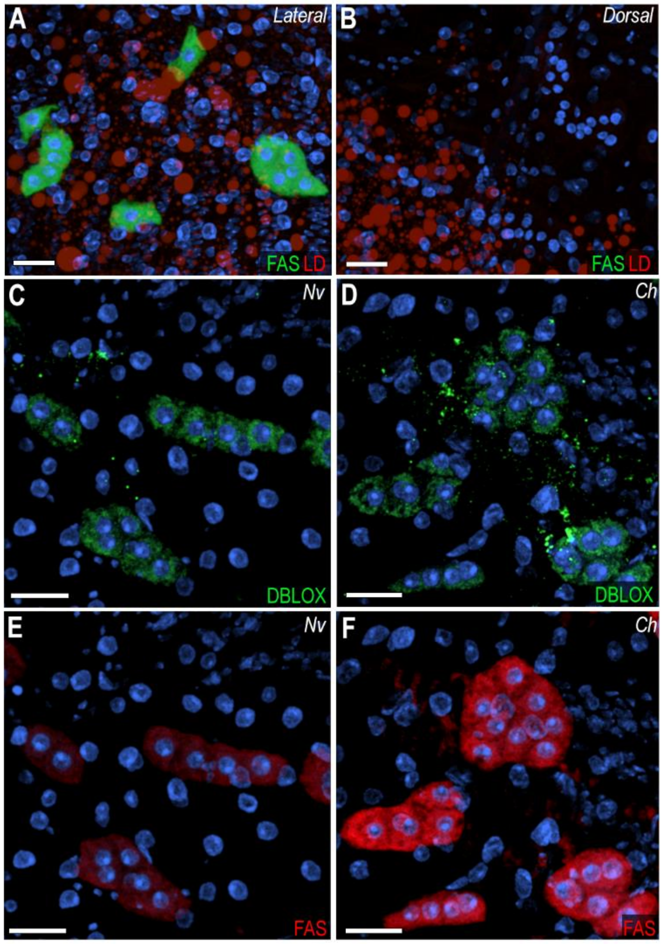


**Supplementary Figure 3. Characterization of oenocytes and their response to immune priming.** (A-B) RNA *in situ* hybridization of Fatty acid synthase (FAS; AGAP001899), a marker of oenocytes, on the lateral (A) and dorsal (B) sides of the abdominal body wall. FAS is shown in green, lipid droplet (LD) in red, and nuclei in blue. (C-H) RNA *in situ* hybridization of FAS and double peroxidase (DBLOX) in ventral abdominal fat body tissue from Naïve (C, E, G) and *Plasmodium berghei*-challenge (D, F,H) conditions. FAS is stained in red, DBLOX in green, and nuclei in blue. Scale bar: 20µm.

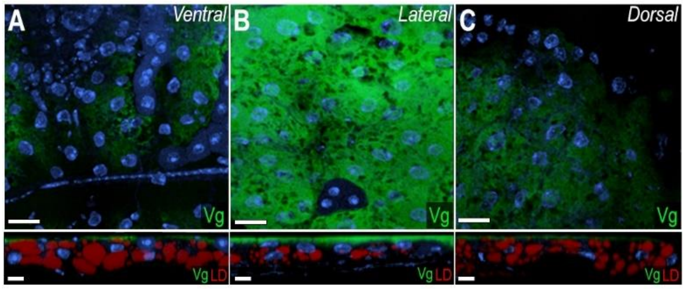


**Supplementary Figure 4. Vitellogenin mRNA expression pattern on abdominal fat body tissue.** (A-C) RNA *in situ* hybridization of Vitellogenin (Vg; AGAP004203) in the abdominal fat body of blood-fed females, shown on the ventral (A), lateral (B), and dorsal (C) sides. (A-C, bottom) Corresponding side views of each region. Vg is shown in green, lipid droplets (LD) in red, and nuclei in blue. Scale bar: 20µm.
